# Tracing the stemness and malignant transition in a heritable colorectal cancer Lynch Syndrome by single-cell RNA-seq analysis

**DOI:** 10.1101/2025.04.06.647495

**Authors:** Junfeng Xu, Yuhang Li, Zhiqin Wang, Qianru Li, Aijun Liu, Jianqiu Sheng, Ge Dong, Lang Yang, Zhigang Cai

**Author notes:** Correspondence (G.D.); (L.Y.); (Z.C.). Junfeng Xu, Yuhang Li, and Zhiqin Wang contributed equally.

## Abstract

**Background:** Lynch Syndrome (LS) is an autosomal dominant disease characterized by germline heterozygous mutations in DNA mismatch repair (MMR) genes. High-risk LS patients may proceed to colorectal cancer (CRC). However, the drivers or biomarkers of LS benign colon tissue approaching malignant CRC are not completely understood. The similarity and difference between LS-related and nonLS-related CRC are also not well interrogated (LS-CRC vs. nonLS-CRC). This study aimed to understand the cellular changes during malignant transition in LS.

**Methods:** Single-cell RNA sequencing (scRNA-seq) was used to analyze paired biopsy samples from 3 patients with LS (cancer tissues vs. adjacent normal tissues, n=3). Single-nuclear RNA sequencing (snRNA-seq) was used to analyze a frozen biopsy sample from a patient with LS (n=1). scRNA-seq or snRNA-seq datasets from CRC were downloaded from the open source. Integrative computational analysis was performed to conclude the distinct pattern in the single-cell atlas. Immuno-histo-fluorescence staining (IHF) were also performed for three key markers.

**Results:** In the single-cell atlas, we observed an increase in primitive cancer stem-cell-like cells with high expression of a general cancer biomarker CarcinoEmbryonic Antigen-related Cell Adhesion Molecule 5 (*CEACAM5*) in the epithelium of the LS. Infiltration of immune cells and DNA repair biological activity are dramatically increased in LS carcinoma. The burden in LS is fundamentally elevated compared to that in CRC or healthy donors when the mutations are in the coding-sequence-wide range. Furthermore, T cell and macrophage-related tumor immunity is readily mobilized in carcinomas compared to paraCArcinomas.

**Conclusions:** This study provides single-cell transcriptomic resource using affected tissues from patients with Lynch Syndrome and describes an integrative profile covering the alterations (cancer stem cell markers, mutation burden, and tumor immunity) during the malignant transition from healthy to Lynch Syndrome and to colorectal cancer at the single-cell level.

**Highlights and Figure/Table Index (Take-Home Messages):**

1. Colon tissues from HD (n=4), patients with LS (n=6) and with CRC (n=3) were collected for snRNA-seq or scRNA-seq analysis; Diagnosis of the 6 LS patients are clearly supported by our pedigree documentations (Fig. 1 and Supp. Fig. 1) and briefed in Table 1;
2. Among the 6 patients with LS, 1 for snRNA-seq analysis (a carcinoma tissue only), 3 for scRNA-seq analysis (3 paired para-carcinoma and carcinoma tissues) and 3 for experimental validations; (Fig. 1)
3. Following studies are focused on the comparisons between HD, LS and CRC; In addition, comparisons between paired carcinoma and para-carcinoma tissues from patients with LS were also performed; (Fig. 2-7)
4. When HD and LS are compared, a cancer stem cell marker *CEACAM5* is readily detected by snRNA-seq analysis; (Fig. 5)
5. When carcinoma and para-carcinoma from LS patients are compared, increased immune cell infiltration of and enhanced DNA repair activity in the malignant tissues is readily detected by scRNA-seq analysis; in addition, three upregulated markers in LS carcinoma (*BACE2*, *GRPC5A* and *OLFM4*) were identified and validated by immunohistoinfloresnce; (Fig. 3 and Fig. 4)
6. SComatic algorithm-based mutation calling analysis suggests a comparable mutation burden between carcinoma and para-carcinoma from LS patients, but such mutation burdens are grossly greater than that from HD or that from patients with CRC; (Fig. 6)

## INTRODUCTION

Human babies carrying germline-dominant heterozygous mutations in one of the DNA mismatch repair (MMR) genes, such as *MLH1*, *MSH2*, *MSH6* and *PMS2* (typically loss-of-function mutations), will develop into Lynch Syndrome (LS) in their middle age (∼40 years old) and further transform into colorectal cancer (CRC) in the later stages of life [1–4]. Because of its close relationship with CRC in the clinic, LS is also known as hereditary nonpolyposis colorectal cancer (HNPCC) according to the Amsterdam criteria, but carriers may also develop extracolonic cancers [1, 5]. It has been reported that in the lifetime of carriers, patients with LS are at very high risk of transforming into CRC (40–60%) or endometrial cancer (40–80%) [3, 6].

**Fig. 1.**
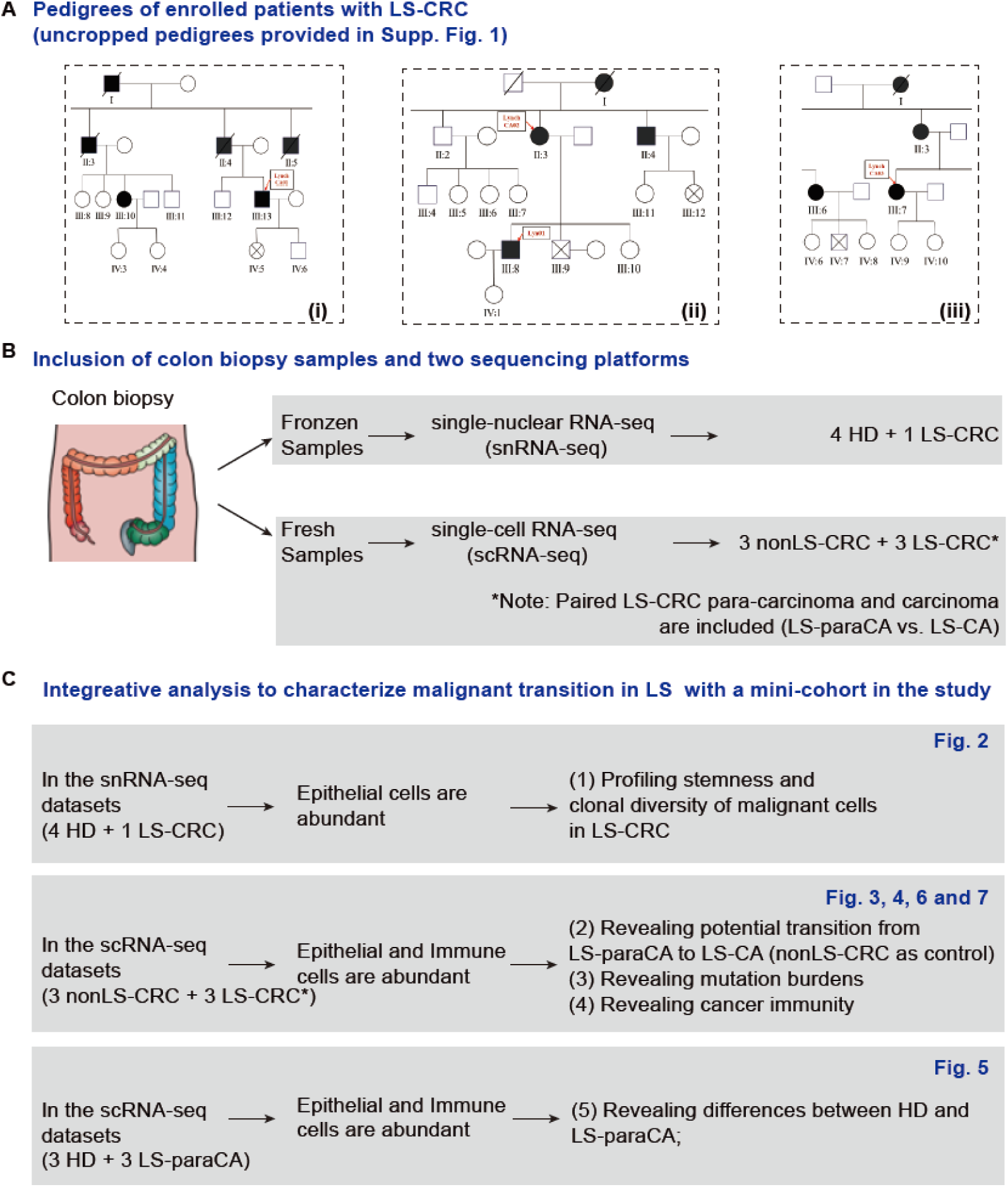
Enrollment of LS patients and overall design of the study. (A) Three family pedigrees of the LS patients (LS-CRC) who donated the samples for single-cell transcriptomic analysis. Cropped pedigree information is provided in the Main text. See the Supp. Fig. 1 for the complete pedigree information. (B) Two different platforms were used in the study: snRNA-seq and scRNA-seq. (C) Various types of comparison were performed to characterize the malignant transition of LS.

**Fig. 2.**
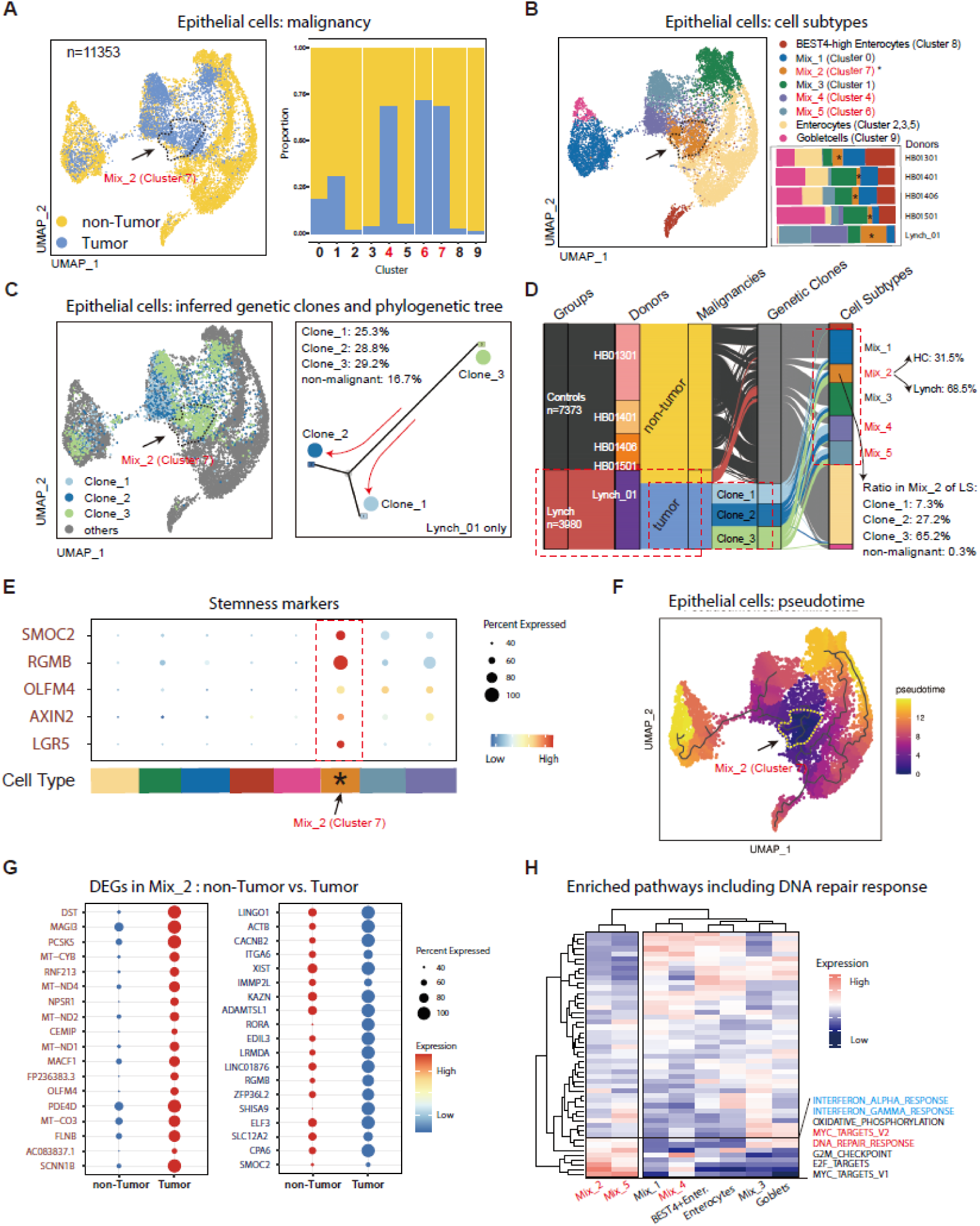
snRNA-seq analysis of a frozen colon biopsy sample from a LS patient reveals a cluster of cancer stem cells. (A) Frozen colon biopsy samples were subjected for snRNA analysis (Healthy donors, N=4; Lynch Syndrome, N=1) and epithelial cells were filtered out for further study. Data were integrated by the Seurat and Harmony standard procedure. Clusters 0-9 were identified in a total of 11,353 cells (the Healthy Controls have 7,373 cells; the LS patient has 3,980 cells). The malignancy of the epithelial cells was inferred by the inferCNV and SCEVAN algorithm (two major compartments: tumor and no-tumor). The portion of tumor or non-tumor cells in each cluster was also illustrated (right panel). Note that the major contributor in Cluster 4, 6, and 7 is the tumor-like cells from the LS patient. (B) Each cluster of the epithelial cells was further annotated according to the combination of Seurat cluster identity and SCEVAN malignant identity. Mix_1 to 5 contain both tumor-like cells and non-tumor-like cells; Other clusters are marked as *BEST4^+^*-Enterocytes, Enterocytes and Goblet cells where much fewer tumor-like cells were assigned. Note that the epithelial cells from the LS patient are mainly classified into Mix_2, 4, and 5 tumor-like cells. (C) Three clones were inferred by SCEVAN algorithm. In the phylogenetic tree, Clone_1 and Clone_2 are closer and probably originated from Clone_3. (D) A Sankey plot showing overall belonging relationships in the epithelial snRNA-seq dataset. Note that the Mix_2 pool is dominated by cells from Lynch_01 and that in the Mix_2 pool Clone_3 is the most dominant clone. (E) The Mix_2 cells specifically express numerous colon epithelial stem cells signature markers such as *SMOC2*, *RGBM*, *OLFM4*, *AXIN2* and *LGR5*. Of note, all these intestine stem cell signature genes were reported as marker genes for cancer stem cells, especially *LGR5*. (F) Pseudotime trajectory analysis of the Lynch Syndrome epithelium cells indicates the Mix_2 cells are the most primitive in development. (G) Top 36 differentially expressed genes (DEGs, both upregulated and downregulated) were highlighted when the non-tumor-like cells were compared with tumor-like cells in the Mix2. (H) The Mix_2, 4, 5 epithelial cells all have enriched signaling pathways involved in DNA repair response and MYC targets (fonts in red) while Mix_4 and 5 manifest Interferon alpha or gamma response (fonts in blue).

**Fig. 3.**
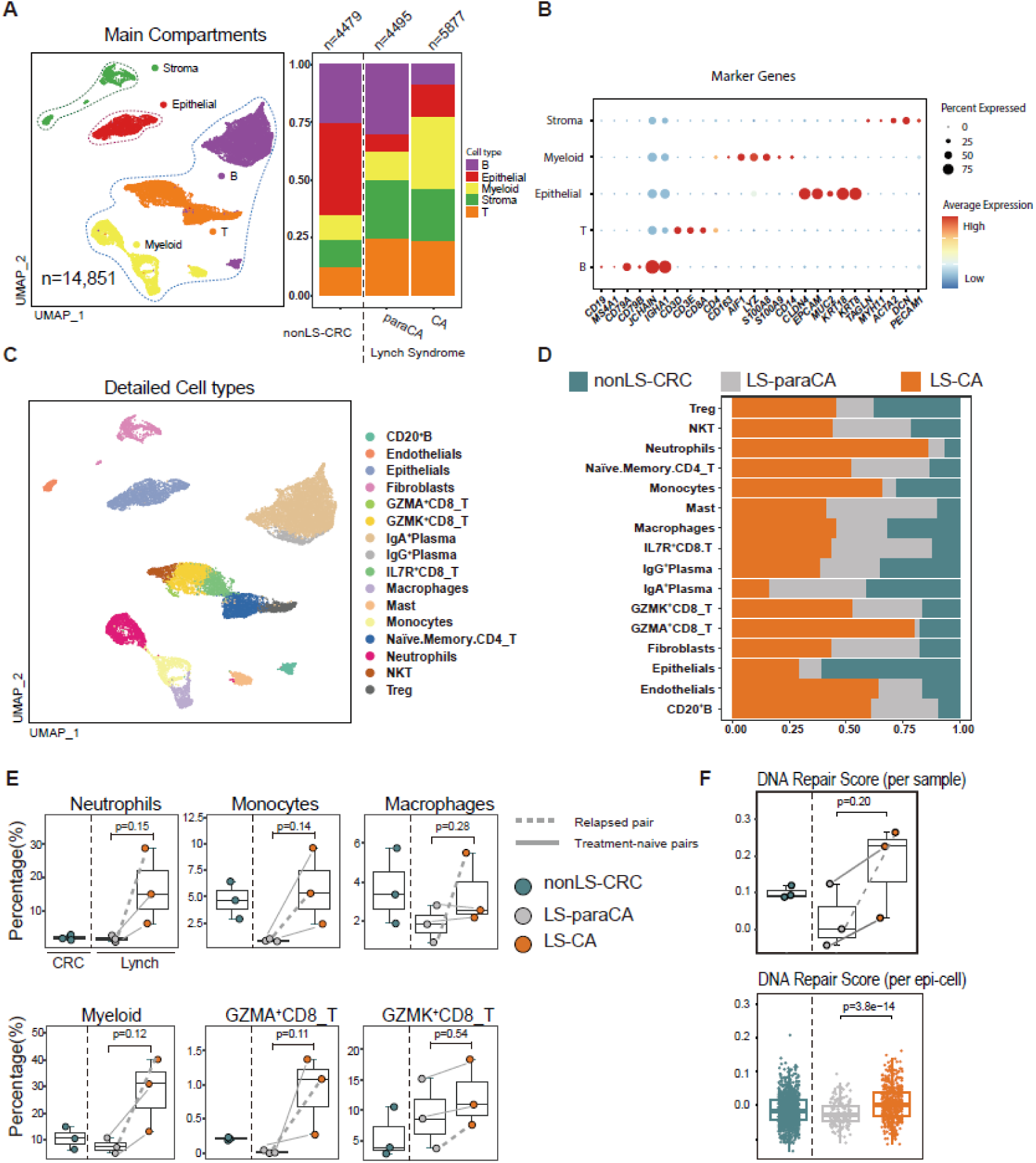
Distinct infiltration of immune cells in carcinoma and para-carcinoma tissues of LS. (A) Fresh paired carcinoma (LS-CA) and adjacent para-carcinoma (LS-paraCA) biopsies of colon tissue was subjected to scRNA-seq analysis (N=3). Colorectal cancer (nonLS-CRC, they are not diagnosed as LS) tumor samples from the public dataset were also included as controls (N=4). Cells were first grossly annotated as stromal, epithelial, or immune cells. Note that the LS-CA group showed increased infiltration of myeloid cells compared with LS-paraCA group, but T cell infiltration was comparable between the two groups. **(B)** Expression of classic markers for annotating the major 5 cell types. **(C)** Detailed annotations of 16 cell types in paired LS-CA and LS-paraCA samples. **(D)** Stacked bar plot showing the contribution of sample sources in each cell type. **(E)** Quantification of immune cell infiltration in LS-CA compared with LS-paraCA. Although no significant differences were observed, the fold-changes were approximately 2–10. Although the overall T cell infiltration was comparable, infiltration of *GZMA*^+^*CD8*_T and *GZMK*^+^*CD8*_T cells appeared to be higher in LS-CA than in LS-paraCA. Of the three LS patients enrolled in the study, one was a relapsed patient (gray dashed line, surgery treatment performed; whose son donated the frozen sample in Figure 2), while the other two were treatment-naïve (gray solid lines). **(F)** Boxplots show DNA repair score quantification per sample (above) or per epi-cell. (bottom).

**Fig. 4.**
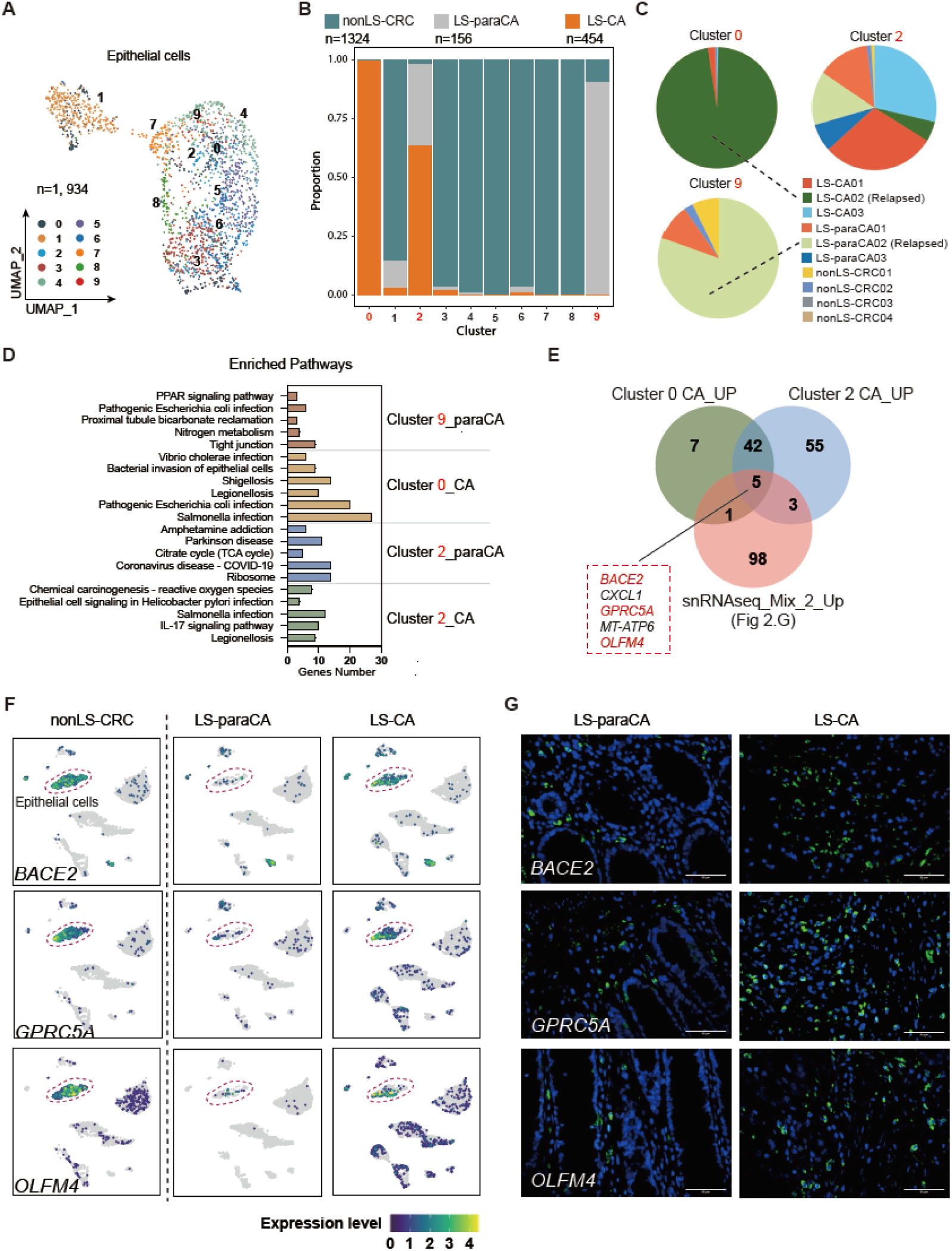
Prioritizing novel epithelial biomarkers for tracing the transition between para-carcinoma and carcinoma in LS (LS-paraCA vs. LS-CA) **(A)** Epithelial cells were filtered out for further analysis and annotated as clusters 0–9 in the UMAP plot. **(B, C)** Bar and pie plots show the contribution of sources to epithelial clusters. Cluster 0 contained cells largely from Lynch CA_02, whereas Cluster 9 was composed of Lynch paraCA_02. The patient who donated Lynch CA_02 and paraCA_02 is identified as a relapsed LS. (D) KEGG enrichment analysis showing the enriched bioprocesses in the four epithelial cell clusters. (E) Five genes with upregulated expression in LS carcinoma epithelial cells were prioritized based on the scRNA-seq and snRNA-seq datasets. (F) Expression of *BACE2, GPRC5A* and *OLFM4* in the paired LS-paraCA and LS-CA groups. Epithelial cells are circled by dashed lines. Note that the expression of genes in epithelial cells from nonLS-CRC samples was also very strong. (G) Verification of the low-to-high transition in the expression of *BACE2, GPRC5A* and *OLFM4*. Paraffin sections from paired LS-paraCA and LS-CA samples were prepared, and immunohistochemical (IHF) staining was performed. The expression of the indicated proteins is labeled in green, whereas the nuclei are stained with DAPI in blue. In total, biopsy samples from 4 patients with LS were used for IHF verification.

**Fig. 5.**
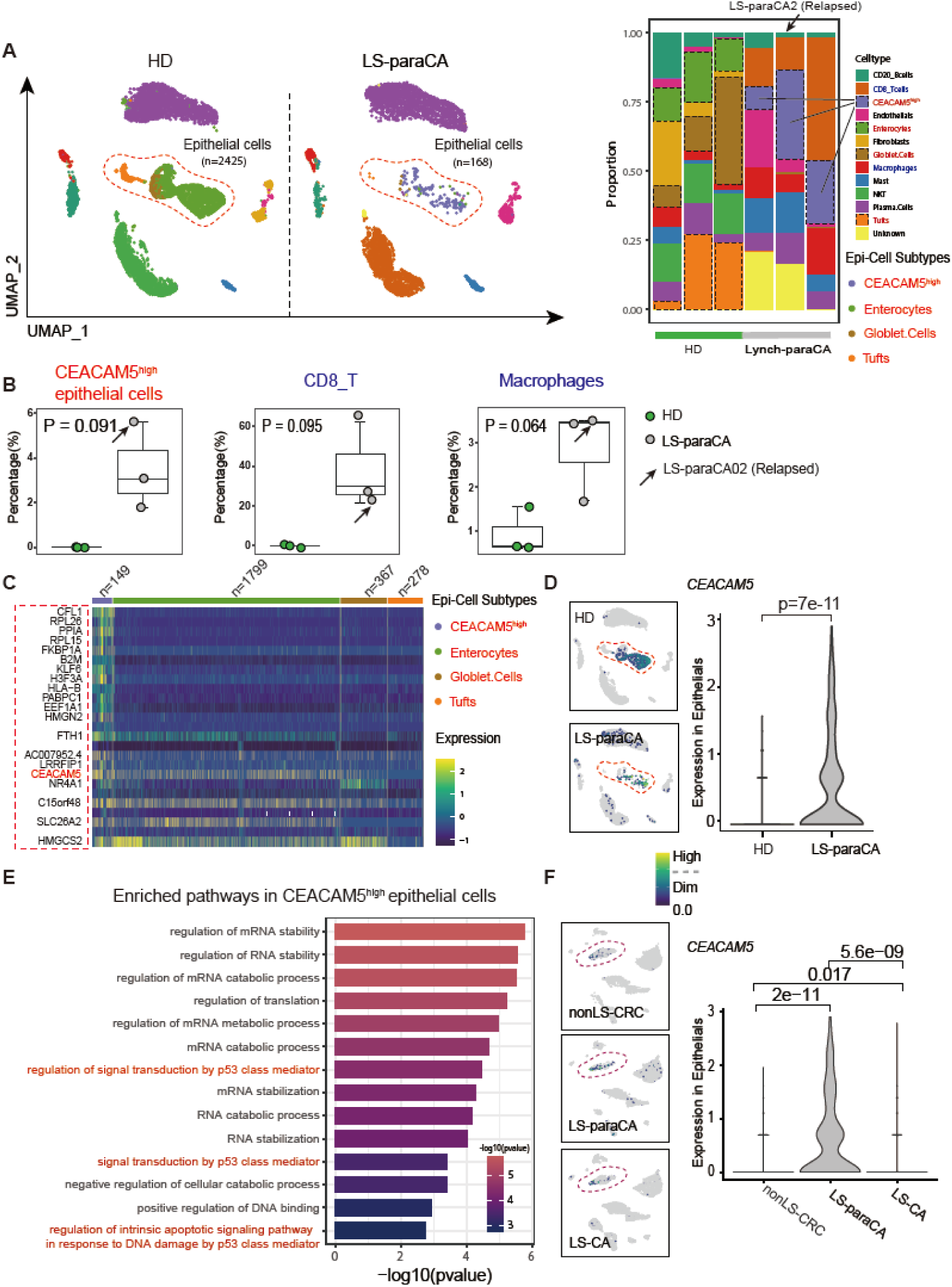
Distinct pre-cancer signs in epithelial and immune cells of LS para-carcinoma. **(A)** The scRNA-seq datasets of LS-paraCA were integrated with those of the colons from healthy donors (HD) (N=3, respectively; the HD group had 7885 cells in total, whereas the LS-paraCA group had 4766 cells). A dominance of *CEACAM5*^high^ epithelial cells were identified in LS-paraCA. The sample from the patient with relapsed LS is marked with an arrow in the bar plot (left panel). **(B)** The portion of *CEACAM5*^high^ epithelial cells, infiltrated *CD8*_T cells and macrophages was quantified. The sample from the relapsed LS patient is marked with arrows. **(C, E)** Top marker genes co-expressed in *CEACAM5*^high^ epithelial cells and the enriched pathways are illustrated in the gene expression heatmap. Cells from the Enterocytes, Tufts, and Goblet subtypes were used as controls. (**D, F)** Quantification of the expression of *CEACAM5* in epithelial cells of HD and LS-paraCA cells.

**Fig. 6.**
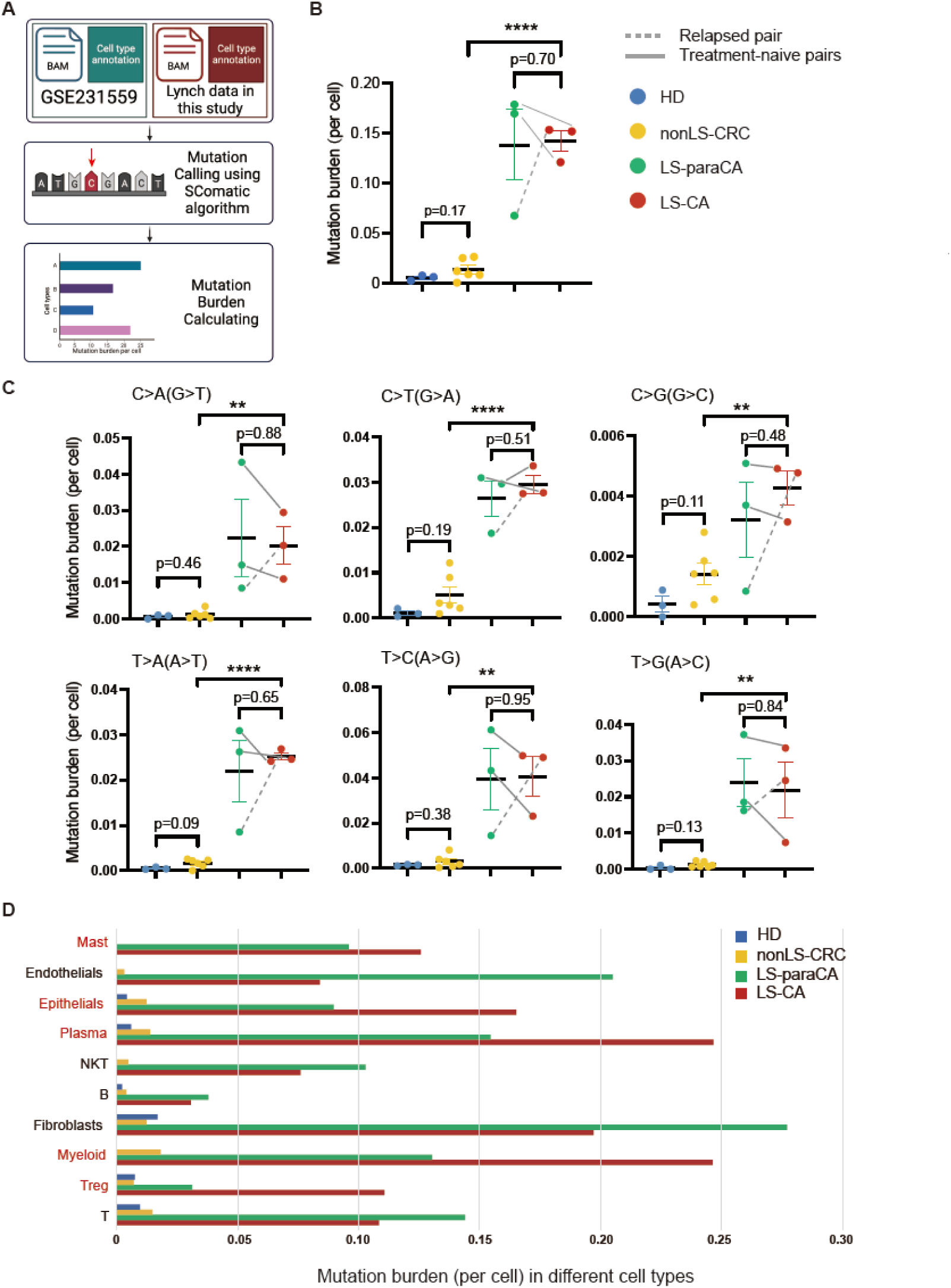
Increased mutation burden in the colon tissues from Lynch Syndrome. **(A)** The schematic diagram for calculating mutation burden based on the scRNA-seq datasets using the SComatic algorithm. Only single-base substitutions (SBS) were called in this study. Once mutation calls were achieved, the mutation burden value was quantified by normalizing the total mutation calls to the total cell number in the samples or cell types (See Methods section for details). **(B)** The total SBS mutation burden in the four groups of samples was indicated. **(C)** The six different SBS mutation burdens in the four groups of samples as indicated. **(D)** The SBS mutation burden in the cell types of the four groups is indicated. Note that the The mutation burden appears to be higher in mast, epithelial, plasma, myeloid, and Treg cells in the LS-CA group than in that the LS-paraCA group.

**Fig. 7.**
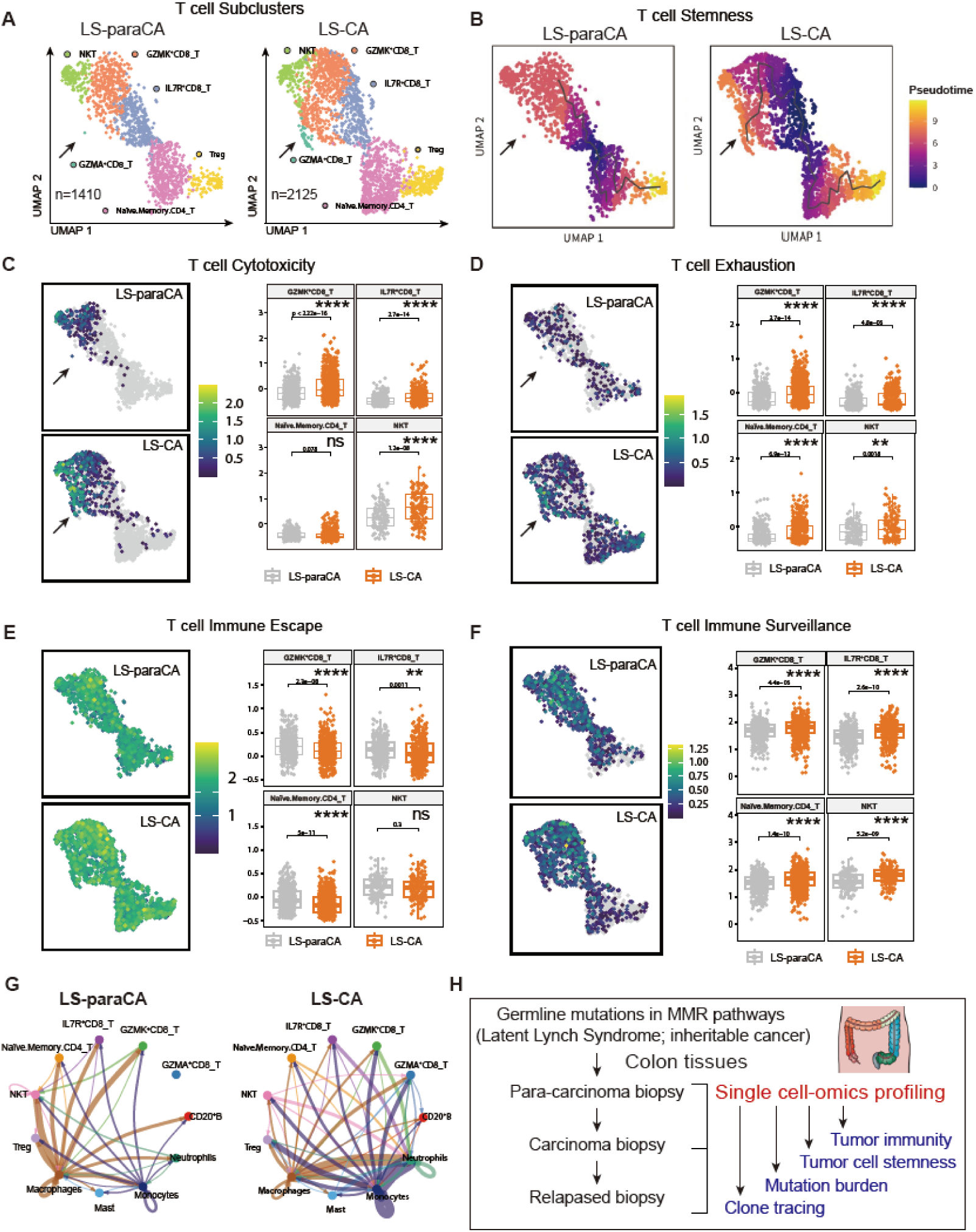
Distinct tumor immunity microenvironment profiles in LS carcinoma. **(A)** T cells from LS-CA and LS-paraCA patients were filtered for further annotation. The LS-paraCA group had 1410 T cells, whereas the Lynch CA group had 2125 T cells. Six major T cell subtypes were identified: Naïve *CD4*-T, NKT, Treg, *IL7R*^+^*CD8*_T, *GZMA*^+^*CD8*_T, and *GZMK*^+^*CD8*_T. Note that *GZMA*^+^*CD8*_T cells appear to be more abundant in Lynch CA (arrows). **(B)** The pseudotime trajectory indicating the T cells developed towards two branches: the Naïve *CD4*-T and Treg branches, and *GZMA*^+^*CD8*_T and *GZMK*^+^*CD8*_T branches. *GZMA*^+^*CD8*_T cells appear to be more mature in LS-CA than in LS-paraCA (arrows). **(C-F)** Biological function-instructed scoring of the paired single-cell datasets of T cells from the three LS patients, including T cell cytotoxicity (**B**), T cell exhaustion (**C**), T cell immune escape (**D**), and T cell surveillance (**E**). **(G)** Circle plots showing interaction profiles across indicated cells in the paired LS samples. The direction of the lines indicates the direction of the cell-cell communication, the width of the lines indicates the interaction strength, and the color of the lines indicates the cell types of the senders. **(H)** Summary of the Study. In this study, we profiled Lynch Syndrome biopsy samples using single-cell or single-nucleus RNA-seq analysis. Paired carcinoma and adjacent para-carcinoma samples were included to facilitate the analysis, including tracing tumor clones, calculating mutation burden, identifying biomarkers for cancer stem cells, and tumor immune microenvironment changes during the transition from pre-carcinoma to carcinoma in the inheritable cancer Lynch Syndrome. See the text in the Discussion section for further detail.

**Table 1.**
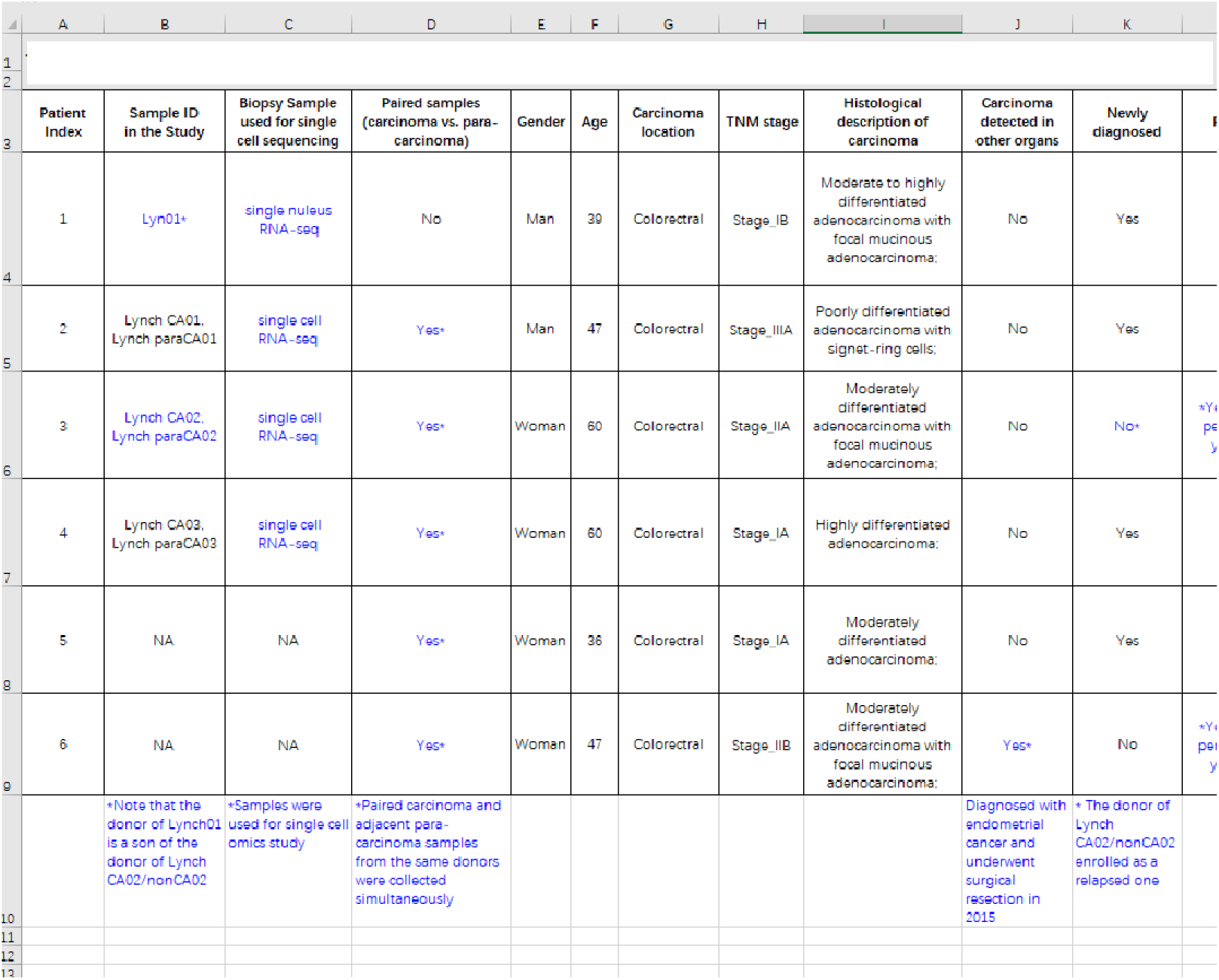
Clinical information of enrolled patients with LS. An Excel version of the table is provided in the Supp. Material

The drivers or biomarkers for malignant transition from latent LS to full-blown LS or CRC during the ages are largely unknown. It has been proposed that malignant onset could be a result of the accumulation of secondary important mutations (also known as the “second-hit” model). For example, mutations in *APC, TP53* or *KRAS* genes, classic drivers of colon cancers, along with mutations in MMR genes, eventually transform LS to CRC [3, 4, 6]. Moreover, except for differences in gene mutation panels for diagnosis, it is not completely understood if LS-CRC is distinct from or similar to nonLS-CRC in the pathology of epithelial carcinoma cells and tumor immunity microenvironment (TIME) [1, 3].

In the last five years, CRC biopsy samples have been broadly examined by single-cell omics in multi-clinical centers, including RNA sequencing, ATAC sequencing, and TCR-sequencing [7–11]. In some studies, CRC patients with mutations in MMR genes were enrolled as sporadic CRC; however, the small subset of CRC patients, LS-CRC, has never been studied in detail with regard to the evolution trajectory of this rare cancer type and any unique features in tumor immunity at the single-cell level [12].

Nevertheless, single-cell sequencing technologies are powerful assets in addition to traditional pathologies and diagnosis. Single-cell omics studies have revolutionized the research of the transition of malignant cells [13]. Even at the most basic level, single-cell sequencing technologies can provide a systematic map (a cell atlas) of tumor progression and alteration of their microenvironment. To specifically profile the cell atlas of LS, a rare heritable cancer, we carried out parallel single-nuclear RNA-sequencing (snRNA-seq) on a frozen LS colon sample and single-cell RNA-sequencing (scRNA-seq) on three paired fresh LS colon samples (carcinoma versus adjacent para-carcinoma). Through a deep bio-computational analysis at both the mutation and functional levels, in this study, we provided a comprehensive single-cell atlas of LS using paired samples and/or family samples. Dramatic increases in immune cell infiltration and DNA repair activity were observed in LS carcinoma compared to the adjacent normal colon tissue. Using healthy and CRC samples as negative and positive controls, respectively, we also observed that LS epithelial cells maintained high stemness activity and had a large portion of cancer-stem-cell-like cells marked by CarcinoEmbryonic Antigen-related Cell Adhesion Molecule 5 (*CEACAM5*). This study highlights important characteristics of the malignant transition of LS.

## MATERIALS and METHODS

### Ethical approval of human sample acquisition, genetic and pathological diagnosis

This study was conducted with the approval of the Seventh Medical Center of the Chinese PLA General Hospital’s Ethics Committee, and participants provided written informed consent prior to participating and in accordance with the 1964 Helsinki Declaration and its later amendments in the study. Patients with LS were enrolled in the hospital between November 2019 and March 2023. The diagnosis of LS was based on the Amsterdam II criteria and germline mutations were subsequently verified. Invasive adenocarcinoma staging was determined based on the analyses of surgically resected specimens using the American Joint Committee on Cancer (AJCC) staging system. Pathologists reviewed counterpart sections of all tumor tissues to confirm the diagnosis. Fresh biopsies were obtained by colonoscopy and single-cell suspensions were immediately prepared and subjected to single-cell RNA sequencing analysis. For single nuclear RNA sequencing, tissues were obtained from patients undergoing surgical resection and frozen at -80 degrees. Five of the six patients with LS donated paired samples (LS CArcinoma and adjacent para-CArcinoma). Clinical information and family relationships of the enrolled patients were summarized in Table 1 (Main text) and Fig. 1A (Main text; See also completed pedigree information provided in Supp. Fig. 1).

### Single-nucleus and Single-cell RNA sequencing

Nuclei were isolated from frozen tissues according to the SeekGene standard protocol. After nuclear dissociation, the cDNA library was constructed using the SeekOne library preparation kit. For fresh samples, after digestion, single-cell suspensions were confirmed with the proportion of living cells exceeding 90% and at a proper concentration of cells greater than 1000 cells/μL. cDNA libraries were constructed within 24 hours. We used the double-ended sequencing mode on the Illumina sequencing platform to perform high-throughput sequencing of the constructed libraries.

### Cell clustering, annotation, and visualization

In-house sequencing data analysis was mainly conducted using Linux-Ubuntu Operation System, R language (version: 4.2), Python language (version: 3.8) and has been described in our previous studies [14, 15]. The R packages includes Seurat, ggPlot2 and In Seurat (version: 4.1), we conducted standard preprocessing and quality control for snRNA-seq and scRNA-seq dataset analyses. We chose the Uniform Manifold Approximation and Projection (UMAP) to perform dimension reduction and visualize the datasets. Harmony (version 0.1) was used to integrate the datasets and control the batch effect. Clusters were identified using the Seurat cluster-finder computation algorithm. Cell types were further annotated based on the expression of canonical tissue compartment markers.

### Malignant cells identification

In snRNA-seq, we extracted all cells sourced from the LS and obtained the raw count matrix. The R package SCEVAN (version: 1.0) was utilized to classify the malignant cells with count matrix input into the core function ‘pipelineCNA,’ which can also be used to infer the copy number profile of malignant cells and identify sub-clone structures [16].

### Pseudotime analysis

Pseudotime trajectory analysis was accomplished by the R package Monocle (version: 3), which was used to predict the evolution trajectory within each time partition and evaluate differentiation levels across cells, based on the expression patterns of key genes. The trajectory roots inferred by Monocle were considered to be at the developmental start points. The cells were colored based on their presumed developmental phases.

### Gene Set Variation Analysis (GSVA)

Gene Set Variation Analysis (GSVA) was used to estimate the variation in pathway activity over a sample population in an unsupervised manner. We conducted ‘GSVA’ and ‘irGSVA’ R packages to calculate enrichment score of the 50 classical hallmark signatures from the GSEA molecular signature database (MsigDB).

### Cell-cell communication analysis

Following Seurat analysis of the scRNA-seq datasets, we further analyzed the differential ligand receptors expressed in different groups with the default parameters, which was accomplished using the R package CellChat (Version: 1.5). The algorithm quantized the number of interactions based on the number of L-R pairs across cells, and the interaction weight was also evaluated by removing the effects of the number of cells.

### Construction of gene signatures and module scoring in each cell

We integrated classic gene sets to characterize different bioprocesses and created gene lists for subsequent analyses. The function ‘AddMouduleScore’ in the Seurat package was used to score the average expression levels for each cluster. All signatures were binned based on the average expression, whereas the control signatures were randomly selected from each bin.

### Immunofluorescence staining

Paraffin-embedded tissue sections were prepared. Tissue sections were dewaxed with xylene, rehydrated, and subjected to antigen retrieval. The slides were blocked with goat serum and incubated with primary antibodies Rabbit monoclonal against BACE2 (1:50, ab270458, Abcam), rabbit polyclonal to GPRCA5 (1:50, ab155557, Abcam), or rabbit polyclonal to OLFM4 (1:50, ab188822, Abcam) at 4 L overnight. Secondary goat anti-rabbit IgG conjugated with Cy5 (1:200; Servicebio) was added and incubated at 37L for 30 min. Nuclei were stained with DAPI. The slides were observed under a laser confocal microscope (Zeiss, Germany).

### Single base substitution (SBS) mutations calling

The biocomputational working platform SComatic is a newly developed tool kit calling somatic mutations in cells, and any germline mutations were filtered out in the procedure [17]. Thus, to our knowledge, SComatic is one of the best algorithms for measuring somatic mutations induced by LS MMR mutations. The SBS calling was conducted in the SComatic software according to the tutorial of SComatic (Python scripts in the Lunix Miniconda environment). First, raw sequencing files (formatted as BAM, containing aligned sequencing reads for all cell types) were split into cell type-specific BAMs using precomputed cell type annotations in the Seurat. Second, the base count information for each cell type and every position in the genome was recorded and merged into a single matrix as TSV files. Finally, variants were called, and high-quality mutations were marked with ‘PASS’ label for downstream statistical analysis. Single base substitutions (SBS) were identified in six types: C>T(G>A), C>G(G>C), C>A(G>T), T>C(A>G), T>C(G>A), T>A(A>T), and T>G(A>C), and the total SBS number is the sum of these six types of SBS. The mutation burden (per sample or cell type) was normalized by dividing the number of mutations by the number of cells in each sample (or cell type).

### Statistical analysis

Statistical analyses were performed using Prism 9 and the R software (version 4.2.1). We chose appropriate statistical methods to calculate two-tailed p-values to evaluate significance. For gene expression, |log_2_ fold change | >0.3 and p<0.05, were considered statistically significant. In other comparative analyses, statistical significance was set at p<0.05.

### Data availability

Healthy and CRC single-cell (or single nucleus) RNA-seq datasets GSE201349, GSE166555, and GSE231559 are publicly available [9, 10, 18]. Lynch Syndrome carcinoma and para-carcinoma scRNA-seq datasets were generated in this study and have been deposited at the China National Center for Bioinformation/Beijing Institute of Genomics, Chinese Academy of Sciences (GSA-Human: xxx): https://ngdc.cncb.ac.cn/gsa-human.

## RESULTS

### Clinical information of the enrolled LS patients

The clinical information of the LS patients (n=6; 2 male and 4 female) enrolled in the study is briefly described in Table 1. A cropped version of the family pedigree is shown in Figure 1A (full version in Supp. Fig. 1). The six patients with LS were from five families (Chinese-Han from 5 different provinces). The years of hospitalization in our center were between 2019 and 2023, and their ages ranged from 36 to 60 years old. All LS biopsy donors met the Amsterdam II criteria and underwent subsequent verification of germline mutations by sequencing a panel of genes, including the four LS-related MMR genes. Germline mutations in genes other than MMR were ruled out. Single-cell sequencing datasets of healthy controls and CRC without indicated LS diagnosis or LS-related MMR somatic mutations (marked as nonLS-CRC hereafter) were downloaded from the public resource. Of the six patients enrolled in the study, two were from a family. The donor of the frozen sample was the son of one of the five paired sample donors, who was diagnosed with a relapsed one (a surgery treatment was performed 8 years ago). One of the six LS patients was diagnosed with endometrial cancer and relapsed LS (surgery treatment was performed 15 years ago), and the other five patients did not report extracolonic cancers. Two different sequencing platforms, snRNA-seq and scRNA-seq, were conducted in the study (Figure 1B). As briefed in Figure 1C, comparisons were designed to characterize the transition between non-malignancies and malignancies in LS (LS-paraCA vs. LS-CA) or between healthy state and pre-malignancies (Healthy vs. LS-paraCA).

### Analysis of a frozen LS colon tissue by snRNA-seq

The frozen malignant colonic tissue from a LS patient (Sample ID in the study: Lynch_01) was used for single nucleus RNA sequencing (snRNA-seq) analysis. Datasets of snRNA-seq of four healthy donors were integrated with Lynch_01 as a control (Sample ID in the study: HB01301, HB01401, HB01406, and HB01501) [10]. After standard quality control, 32,083 nuclear transcriptomes were used for downstream analyses. Three major cell types, epithelial cells, stromal cells, and immune cells, were annotated. The Lynch_01 sample exhibited an increase in epithelial cells compared to controls but had much fewer stromal and immune cells in the frozen sample (in Lynch_01, the percentage of epithelial cells was 95.3%, while that of stromal and immune cells was 3.6% and 1.1%, respectively).

To characterize the alterations in the frozen LS samples, only epithelial cells were used for further downstream analysis (Figure 2; the total epithelial cell number was 11353 and Lynch_01 had 3980 cells). As shown in Figure 2A-D, clusters 0 to 9 were annotated by the standard Seurat analysis, while tumor and non-tumor cells were inferred from the epithelial cell pool based on copy number variants (CNVs) using the SCEVAN algorithm [16]. Epithelial clusters 2, 3, 5, 8, and 9 had relatively low proportions of tumor-like cells. However, clusters 0, 1, 4, 6, and 7 exhibited a high proportion of tumor-like cells, and these clusters were renamed Mix_1 to 5, as indicated (Figure 2B). Of note, most of the cells in Lynch_01 were recognized as tumor-like cells (tumor cells: 83.3%; non-tumor cells: 16.7%), and three clones and their phylogenetic tree are indicated (Figure 2C). Although the phylogenetic tree suggested that clones 1 and 2 were more correlated, the ratios of the three clones in the epithelium and their flow in the epithelial cell subtypes were comparable, suggesting that no distinct bias between the clones was observed in the development of the eight epithelial subtype cells (Mix 1 to 5, *BEST4*^high^ enterocytes, enterocytes, and goblet cells; Figure 2D).

Interestingly, the expression of numerous markers and pseudotime analysis suggested that the pool of Mix_2 epithelial cells (Cluster 7) was dominated by intestinal stem cells (Figure 2E, F). Five gene signatures, including the well-known *LGR5,* are also recognized as intestinal cancer stem cell marker genes [19–23]. Quantification of Mix_2 in the samples suggested that Lynch_01 had a much higher proportion of Mix_2 epithelial stem cells than the four healthy controls (16.2% in Lynch_01 vs. 4.0% in HC). Furthermore, almost all Lynch_01 cells in Mix_2 were inferred as malignant cells (tumor-like cells; 643 cells out of 645 cells). The proportion of Clone_3 in Mix_2 was the highest (Clone_3,65.2%; Clone_2,27.2%; Clone_3,7.3%; non-malignant, 0.3%) (Figure 2D). Differential gene expression analysis in the Mix_2 pool revealed that genes, including *DST* and *MAGI*, were upregulated in the tumor-like sub-pool, while genes, including *LINGO1* and *ACTB*, were downregulated in the tumor-like subgroup compared to those in the non-tumor-like sub-pool (Figure 2G). Accordingly, we also revealed that Mix_2, 4, and 5 pools (dominated by the tumor-like cells from Lynch_01) exhibit much higher activity in DNA repair and MYC_ target pathways (Figure 2H), strongly suggesting the MMR pathway and malignant activity in Lynch_01 compared to healthy controls.

### A cell atlas of LS carcinoma and para-carcinoma revealed by single cell sequencing

To overcome the limitations posed by the absence of immune cells in snRNA-seq analysis, we conducted single-cell RNA sequencing analysis (scRNA-seq) using paired fresh carcinoma samples and their adjacent tissues from three LS patients. Datasets of three randomly selected CRC patients were included as malignant controls (GSE166555) [9]. As shown in Figure 3A, both LS carcinoma and para-carcinoma samples had major cell types, including stromal cells, epithelial cells, and infiltrated immune cells (B cells, myeloid cells, and T cells). The expression of the classic markers is shown in Figure 3B. The cells were further annotated into 16 cell types, as shown in Figure 3C. The portion between sample groups is shown in Figure 3D. Intestinally, increased infiltration of many types of immune cells was observed in LS carcinoma (LS-CA) compared with para-carcinoma (LS-paraCA) (LS-CA vs. LS-paraCA) (Figure 3E). Of note, among the three LS patients, the relapsed donor always exhibited a pattern of increased infiltration of neutrophils, monocytes, macrophages, *GZMA*^+^*CD8*_T, and *GZMK*^+^*CD8*_T, suggesting that LS carcinoma has an increased immune response compared to latent LS-paraCA tissue (dashed lines, Figure 3E). Accordingly, when DNA repair activity was measured in the cell atlas, we observed that the activity was increased in LS-CA compared to that in LS-paraCA (Figure 3F). These results demonstrate that the study provides for the first time a cell atlas for comparing LS carcinoma and para-carcinoma, and in the atlas, infiltration of immune cells and DNA repair activity are readily detected at a higher level in LS carcinoma than in para-carcinoma.

### Prioritizing novel biomarkers tracing the malignant transition in the paired LS epithelial cells

Although there were not enough epithelial cells available to perform similar clonal analysis by SCVAN in each LS paired sample, we were able to filter out epithelial cells and compare the expression of important genes between LS-CA and LS-paraCA. Seurat analysis of the epithelial cells revealed that 10 different clusters were recognized and that Clusters 0, 2, and 9 were dominated by LS samples (Figure 4A-C). We enriched the biological pathways in these clusters using KEGG analysis (Figure 4D). Furthermore, we prioritized five conserved genes that were upregulated in Mix_2 of snRNA-seq (Figure 2G) and in Cluster_0_CA and Cluster_2_CA: *BACE2*, *CXCL1*, *GPRC5A*, *MT-ATP6* and *OLFM4* (Figure 4E). As *CXCL1* encodes a secreted protein and *MT-ATP6* encodes a mitochondrial protein, we chose the other three genes to verify whether their expression is dominant in epithelial cells and to perform protein-level verification using immunohistofluorescence (IHF) experiments. As shown in Figures 4F and G, the expression of *BACE2*, *GPRC5A* and *OLFM4* and their encoded proteins was much higher in malignant CRC and LS-CA tissues, while their expression in LS-paraCA tissues was limited. In summary, these results demonstrate that LS patients have a unique epithelial cell cluster distinct from CRC tissue and that a series of conserved biomarkers should be used to trace malignant transition in LS.

### *CEACAM5* as an early biomarker for abnormal epithelial cells in latent LS samples

LS is a disease induced by germline mutations, and the malignant sign is probably much earlier than the appearance of carcinoma tissue. This suggests that the epithelial cells from LS-paraCA patients may show some abnormalities when compared to healthy controls. We then integrated our datasets of LS-paraCA epithelial cells with those of three healthy donors (GSE166555) to determine whether markers earlier than the above three biomarkers exist [9]. As shown in Figure 5A to C, a *CEACAM5*^high^ epithelial cell cluster was recognized based on the single-cell atlas, and it appears that such a cluster is specific to LS patients. *CEACAM5* is well-known cancer stem cell biomarker and immune checkpoint target [24, 25]. Interestingly, the proportion of *CD8* + T cells and macrophages, two important immune cells in the tumor microenvironment, was much higher in the LS-paraCA group than in the healthy controls (HD) (Figure 5C). The genes that were highly co-expressed with *CEACAM5* were plotted in a gene expression heatmap, and the enriched pathways included RNA stability, metabolic-related processes, and *p53*-related DNA damage pathways (Figure 5C and E). As *CEACAM5* expression is not specific to *CEACAM5*^high^ epithelial cells and LS patients, we quantified the level of *CEACAM5* in healthy controls, LS-paraCA, LS-CA, and nonLS-CRC patients as controls. As shown in Figure 5D, healthy controls only maintained a very dim expression level of *CEACAM5* in epithelial cells, while LS-paraCA manifested a much higher expression of that. Similarly, *CEACAM5* was expressed in nonLS-CRC and LS-CA cells (Figure 5F). Taken together, these results suggest that LS-paraCA may confer biomarkers such as *CEACAM5* and infiltration of immune cells much earlier than previously thought. These biomarkers should be tracked along with LS progression as early as possible to monitor malignant transformation in patients’ management.

### Comparable high SBS mutation burden in LS carcinoma and para-carcinoma at the coding-sequence-wide level

The “second-hit” model has been proposed to explain the malignant transition in the LS, indicating LS carcinoma tissue may have an increased tumor mutation burden than that in the LS para-carcinoma tissues. To test if this hypothesis is true, we measured the single-base substitution (SBS) mutation burden at the coding-sequence wide (rather than genome-wide) using the SComatic algorithm tool. The computational analysis procedure is illustrated in Figure 6A (see reference for details) [17]. To test whether any differences in SBS burden exist between LS and non-LS CRC, datasets of three healthy controls and six CRC patients were also included (GSE231559) [18]. As shown in Figure 6A, the SBS mutation burden was greater than 10 folds higher in LS-CA and LS-paraCA than in that in nonLS-CRC (average fold change: 10.7, p<0.001). As expected, the mutation burden in LS-paraCA CRC patients was also much higher than that in healthy controls (fold change on average: 1.7, p=0.17). LS-CA and LS-paraCA maintained a high SBS mutation burden; however, no significant differences were detected between LS-CA and LS-paraCA at the total level or the level of each single SBS pattern (fold change in average: 1.1, p=0.7, Figure 6B). We also measured the SBS mutation burden in the six main types, and no significant differences were observed (Figure 6C). Notably, the relapsed LS patient always exhibited an SBS mutation burden in the CA tissue compared to that in the paraCA tissue (Figure 6B, C). Furthermore, mast, epithelial, plasma, myeloid, and T cells appeared to have a higher mutation burden in LS-CA than that in LS-paraCA (Figure 6D). In summary, these results demonstrate that LS maintains a comparably high level of SBS mutation burden, but no significant difference was detected between carcinoma and para-carcinoma tissues.

### Tumor immunity is mobilized during the transition from latent LS to full-blown LS

The cell atlas of LS also facilitates detailed profiling of alterations in the TIME of heritable cancer. As shown in Figures 3E and 5B, we demonstrated that immune cells are dramatically mobilized when comparing LS-CA and LS-paraCA or comparing LS-paraCA and healthy controls (HD). To dissect the molecular features of tumor immunity in detail in patients with LS, we isolated T cells for further analyses (Figure 7A). Pseudotime trajectory analysis revealed T cells branching into *CD4* and *CD8* categories, with Naive Memory *CD4*^+^T cells evolving into Treg cells, and *IL7R*^high^*CD8*^+^T cells transitioning into *GZMK*^high^*CD8*^+^T, *GZMA*^high^*CD8*^+^T, and NKT cells (Figure 7B). Functional assessment of T cells using established canonical gene sets indicated consistent trends in LS carcinoma (LS-CA vs. LS-paraCA), with enhanced cytotoxicity, exhaustion, and immune surveillance across T cell subtypes (Figure 7C-F). Furthermore, the overall profiles of cell-cell interaction strength were more complex and abundant in LS-CA than in that in LS-paraCA (Figure 7G). In summary, these results demonstrate that immune cells were more actively mobilized into the malignant tissue in LS and that immune cell-cell communication was also more active in malignant LS tissue [26].

## DISCUSSION

In this study, we focused on a specific group of colorectal cancer patients diagnosed with Lynch Syndrome and profiled the malignant transition using single-cell sequencing and single-nuclear sequencing.

As outlined in Figure 7H, our study demonstrated that: 1) both frozen samples and fresh samples are feasible for single-cell (or nucleus) analysis (median UMI numbers are between 1120 and 2638 with snRNA-seq having the lowest UMI); 2) DNA repair activity is readily detected at the gene expression level when epithelial cells of LS are compared with normal epithelial cells; 3) SBS mutation burden is comparable in LS-CA and LS-paraCA but greater than that in CRC or healthy donors; 4) LS carcinoma has a high stemness score and numerous biomarkers including *CEACAM5*, *BACE2*, *GPRC5A* and *OLFM4* may be utilized as novel molecular markers for tracing the malignant transition in LS. 5) The infiltration and activity of immune cells are greatly mobilized during the transition from a non-malignant state to a malignant state in the LS. This study set up a comprehensive computational workflow for observing neoplasm transition in the real world for LS by utilizing a single-cell sequencing dataset.

### Limitations of the study

#### I. Limitations of acquired clinical samples

An obvious limitation of the present study is that only four LS patients were enrolled for a single-cell omics study plus an additional two for experimental verification, and no longitudinal biopsy samples were included. Considering that clinical studies rely on biopsy sample availability in practice, the six donors represent a small LS population. Nonetheless, this study is certainly instructive to the field since we are using single-cell sequencing technologies to monitor the transition (or the difference) between latent LS colon tissue and diseased (malignant) LS colon tissue. Importantly, two members of a family with LS were enrolled in the study (a mother and a son), and the mother was a relapsed LS patient, while the other three LS patients were treatment-naive. This diversified representation of the enrolled LS cohort with the scRNA-seq dataset represents an invaluable resource for future LS studies.

Another obvious limitation of the present study is that no whole-exon-wide sequencing (WES) analysis was conducted to validate the findings, as we only used the availability of the scRNA-seq dataset to compare the mutation burden of LS-CA and LS-paraCA [27]. Reutilization of the scRNA-seq dataset by the SComatic algorithm enabled us to compute the SBS mutation burden without additional sequencing costs. The addition of WES sequencing to paired samples of colon tissue or single-cell DNA sequencing offers an opportunity to observe the mutation burden at the whole-genome and single-cell levels.

#### II. Limitations of our bio-computation analysis and the future direction of the study

The present study largely relies on the computational analysis of scRNA-seq or snRNA-seq datasets with transcriptomic features and a large number of cells. Annotation of the single cells with a regular analysis protocol on the Seurat platform enabled us to identify the changes in the proportion of cell types and cell-cell communications. Using SCVAN and SComatic algorithms, we were also able to profile CNV alterations and SBS mutations. These two algorithms are particularly important because LS is a heritable CRC with mutations in MMR genes. Furthermore, we validated our findings using immunohistochemistry and flow cytometry experiments. However, full-length sequencing of the cDNA will be more informative for the SComatic analysis. Combination of scRNA-seq and scATAC-seq will assist the cell atlas study of LS as well. We envision that an improved bio-computational analysis working-flow will be instructive for future LS studies when more donors are enrolled.

## CONCLUSION

Taken together, this study provides a cellular atlas of LS colon diseased tissues at a single-cell level. Through paired samples, we profiled the alterations in cancer stem cell markers and infiltrated tumor immune cells during the transition of latent LS and full-blown LS. Using a limited number of enrolled patients, we did not observe a dramatic difference in SBS during the transition; however, an increase in DNA repair activity was detected. This study suggests several novel cancer stem cell biomarkers for monitoring LS progression and suggests that single-cell sequencing technologies will empower a deeper understanding of the pathologies of LS, the heritable colorectal cancer.

## Supp. Files

**Supp. Fig. 1 Uncropped version of family pedigree of patients with Lynch syndrome.** Related to Figure 1A.

(A) Family pedigree of Lynch CA01.

(B) Family pedigree of Lynch CA02 and Lyn01.

(C) Family pedigree of Lynch CA03.

All cases included in the study met the Amsterdam II criteria: 1) three relatives with Lynch Syndrome-related cancers; 2) one of which is a first-degree relative of the other two; 3) Lynch Syndrome-related cancer affects more than one generation; and 4) at least one Lynch syndrome-related cancer diagnosed before the age of 50 years. (a Lynch Syndrome-related cancer sites included the colon, rectum, endometrium, ovary (including fallopian), sebaceous carcinoma, small bowel, ureteric, or CNS gliomas (including glioblastoma and astrocytoma).

## Abbreviations

LS, Lynch syndrome; MMR: Mismatch repair; CRC: Colorectal cancer; CEACAM5: Carcinoembryonic antigen-related cell adhesion molecule 5; SBS, single-base substitution; HNPCC: Hereditary nonpolyposis colorectal cancer; TIME: Tumor immunity microenvironment; GSVA: Gene set variation analysis; CNVs, copy number variants; WES, whole-exon-wide sequencing.

## Acknowledgments

We thank our colleagues for their technical support, critical reading of our manuscript, and suggestions for improving the manuscript.

## Authors’ contributions

Junfeng Xu, Ph.D., M.D. (Conceptualization: Equal; Data curation: Equal; Formal analysis: equal; investigation: equal; methodology: equal; resources: equal; validation: lead; visualization: equal; writing the original draft: equal).

Yuhang Li, Ph.D. (Conceptualization: Equal; Data curation: Equal; Formal analysis: lead; methodology: equal; visualization: equal; writing the original draft: equal).

Zhiqin Wang, Ph.D. (Conceptualization: Equal; Data curation: Equal; Formal analysis: Lead; Methodology: Equal; Visualization: Equal; Writing-original draft: Equal; Writing-review and editing: Supporting)

Qianru Li, Ph.D. (Formal analysis: Supporting; Methodology: Supporting)

Aijun Liu, Ph.D., M.D. (Data curation: Supporting; Formal analysis: Supporting)

Jianqiu Sheng, Ph.D., M.D. (Funding acquisition: Lead; Project administration: Supporting; Supervision: Supporting; Writing-review and editing: Supporting)

Ge Dong, Ph.D. (Writing-review & editing: Supporting)

Lang Yang, Ph.D., M.D. (Conceptualization: Lead; Formal analysis: support; funding acquisition: Lead; Project administration: support; writing: review and editing: lead).

Zhigang Cai, Ph.D. (Conceptualization: Lead; Methodology: Lead; Funding acquisition: Equal; Resources: Lead; Supervision: Lead; Project administration: lead; visualization: lead; and review and editing: lead).

## Data Availability

The sequencing data in this study were deposited at the China National Center for Bioinformation/Beijing Institute of Genomics, Chinese Academy of Sciences (GSA-Human: XXX): https://ngdc.cncb.ac.cn/gsa-human.

